# Deletion of *Pax1* scoliosis-associated regulatory elements leads to a female-biased tail abnormality

**DOI:** 10.1101/2023.04.12.536497

**Authors:** Aki Ushiki, Rory R. Sheng, Yichi Zhang, Jingjing Zhao, Mai Nobuhara, Elizabeth Murray, Xin Ruan, Jonathan J. Rios, Carol A. Wise, Nadav Ahituv

**Author notes:** These individuals contributed equally to the study (Aki Ushiki, Rory R. Sheng).

## Abstract

Adolescent idiopathic scoliosis (AIS), a sideways curvature of the spine, is sexually dimorphic, with increased incidence in females. A GWAS identified a female-specific AIS susceptibility locus near the *PAX1* gene. Here, we used mouse enhancer assays, three mouse enhancer knockouts and subsequent phenotypic analyses to characterize this region. Using mouse enhancer assays, we characterized a sequence, PEC7, that overlaps the AIS-associated variant, and found it to be active in the tail tip and intervertebral disc. Removal of PEC7 or Xe1, a known sclerotome enhancer nearby, and deletion of both sequences led to a kinky phenotype only in the Xe1 and combined (Xe1+PEC7) knockouts, with only the latter showing a female sex dimorphic phenotype. Extensive phenotypic characterization of these mouse lines implicated several differentially expressed genes and estrogen signaling in the sex dimorphic bias. In summary, our work functionally characterizes an AIS-associated locus and dissects the mechanism for its sexual dimorphism.

## Introduction

Adolescent idiopathic scoliosis (AIS), characterized by a lateral curvature of the spine that occurs during early adolescence along with spine growth, affects ∼3% of the population worldwide^1^. AIS is a sexually dimorphic disease^2^. Girls have approximately ten-fold higher risk of developing progressive curves that require operative treatment^3^. AIS is caused by both environmental and genetic factors. A study of monozygotic and dizygotic twins from the Swedish twin registry estimated that the overall genetic effects for AIS accounted for 38% of the observed phenotypic variance^4^. However, the genetic basis and pathogenic mechanism of AIS have remained largely unknown. Genome-wide association studies (GWAS) have identified more than a dozen genetic loci associated with AIS with the majority located in noncoding regions of the genome^5–14^. These AIS-associated single nucleotide polymorphisms (SNPs) potentially overlap gene regulatory sequences, such as enhancers, and could alter their target gene expression^15^.

In one of these GWAS, we identified an AIS susceptibility locus that was associated only with females, downstream of the paired box 1 transcription factor (*PAX1*), a transcription factor involved in sclerotome development^10^. *Pax1* is expressed in the developing spine in mouse embryos. Both heterozygous and homozygous *Pax1* knockout (KO) mice exhibit a kinky tail phenotype^16,17^, and various spinal malformations, including scoliosis^18,19^. PAX1 regulates extracellular matrix (ECM) genes, such as collagen and aggrecan, and is crucial for mesenchyme condensation and intervertebral disc (IVD) development^20,21^. Despite the growing understanding of the function of PAX1 in spine development, how it leads to AIS remains largely unknown.

Several *Pax1* associated gene regulatory elements have been characterized. One such element, Xe1, which resides near the AIS-associated GWAS SNP (**Fig.1a**), was found to drive enhancer activity in the mouse sclerotome^22^. Additionally, using zebrafish enhancer assays, our lab functionally characterized ten enhancer candidate sequences in the *PAX1* AIS-associated locus, termed *PAX1* Enhancer Candidates (PECs)^10^. These PECs were chosen based on their evolutionary conservation and/or having enhancer-associated marks in ENCODE datasets^23^. Only two of these sequences drove enhancer activity in the developing zebrafish spine and somitic muscle, the previously characterized Xe1 and PEC7, which is in close vicinity (1,037 base pairs (bp)) to Xe1 (**Fig.1a**). Of the two, only PEC7 harbors an AIS GWAS associated variant, rs169311, which is in strong linkage disequilibrium with the AIS lead SNP, rs6137473. Introducing the AIS-associated SNP rs169311 into PEC7 abolishes its enhancer activity in zebrafish^10^, suggesting that rs169311 downregulates *PAX1* gene expression in the developing somites. However, the mechanisms of how this SNP alters PEC7 function and how this altered function leads to AIS have not been characterized.

**Fig. 1.**
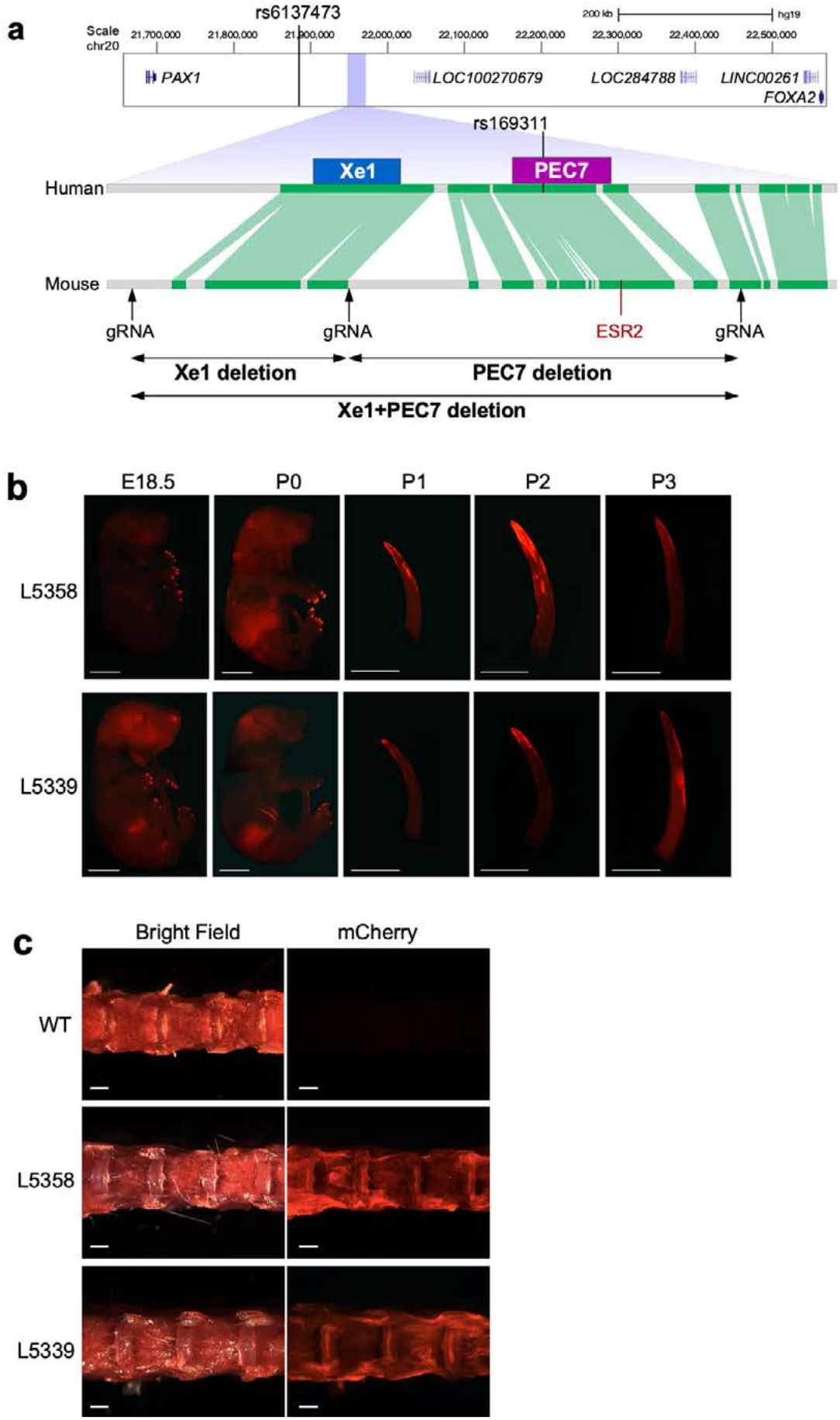
*PAX1* genomic locus in humans and mice and PEC7 enhancer transgenic assay. **a**, Comparison of the *PAX1* locus between human and mouse. The conservation track is from the Ensembl Genome Browser with green lines indicating conserved sequences between humans and mice. The location of the AIS-associated SNPs and gRNAs are also shown. **b**, mCherry expression in PEC7-HSP68-mCherry transgenic mice from E18.5 to P3 (lines 5358 and 5339). White bar represents 5 mm. **c**, mCherry expression in lumber IVD of 10-week-old mouse. White bars represent 1 mm.

In this study, we further characterized the role of this female AIS-associated region using multiple mouse models. Using transgenic mouse enhancer assays, we find PEC7 to be active in neonate tail tip region and adult intervertebral disc (IVD). We next generated three putative *Pax1* enhancer knockout mouse lines— Xe1, PEC7, and both Xe1 and PEC7 (Xe1+PEC7), using CRISPR/Cas9 genome editing. We found that approximately 20% of Xe1 homozygous KO mice have a kinky tail phenotype. This phenotype becomes more severe in Xe1+PEC7 mice, and, intriguingly, it has a higher penetrance in Xe1+PEC7 KO females (60%) than males (42%). To characterize the mechanism leading to the kinky tail phenotype, we performed RNA-seq on embryonic day 12.5 (E12.5) tails, finding ECM genes, including collagen- and aggrecan-encoding genes, to be downregulated in all three homozygous knockout mice. To further characterize the observed sexual dimorphic phenotype, we injected tamoxifen, an estrogen receptor antagonist, to E17.5 Xe1+PEC7 homozygous pregnant females and found that the kinky tail sex ratios was reduced to similar levels as the Xe1 knockout mice (25% in males and 27% in females), suggesting that estrogen could be involved in this sex bias. RNA-seq analysis of postnatal day 2 (P2) tail samples of tamoxifen injected versus un-injected Xe1+PEC7 mice identified genes involved in the reversal of this sexual dimorphic kinky tail phenotype. Combined, our results show that PEC7 functions as an enhancer in the neonate tail tip region and adult IVD and contributes to the onset and progression of a sex-differential kinky tail phenotype, likely due to the estrogen signaling pathway.

## Results

### PEC7 shows enhancer activity in the distal part of the neonatal tail and adult IVD

We first set out to test the mouse enhancer activity of PEC7, as it encompasses rs169311, an AIS GWAS associated variant that abolished its somite enhancer activity when tested in zebrafish^10,24^. We cloned mouse PEC7 into the Hsp68-LacZ vector, in front of an *Hsp68* minimal promoter followed by mCherry reporter gene^25^. This plasmid was injected into mouse zygotes, and two independent transgenic mouse lines were established (line 5339 and line 5358). We checked whole-mount mCherry expression at E12.5, E15.5, E18.5, and P0 to P4, and observed comparable enhancer activity in both lines (**Fig. 1b, Supplementary Fig. 1a**). From E18.5, mCherry was observed in the nails, followed by the distal part of the tail in neonatal stages (P0-P3), with P2 showing the highest mCherry expression which gradually decreased in P3 and P4 (**Fig. 1b, Supplementary Fig. 1a**).

These results suggest that PEC7 has enhancer activity in the distal part of the neonate tail and adult IVD.

### Xe1 + PEC7 homozygous mice have a kinky tail with altered sex penetrance

To characterize the function of this female associated AIS locus, we generated various mouse knockouts for this region. PEC7 is located nearby Xe1 (1 kb in human and 4.3 kb in FVB mice; **Fig. 1a**), a previously characterized sclerotome enhancer^22^, and might be associated with its regulation. Therefore, we generated three different deletion lines: 1) PEC7 only; 2) Xe1 only; 3) PEC7 and Xe1. We first identified the conserved region between humans and mice using the Ensemble genome browser^24^, finding both Xe1 and PEC7 to be sufficiently conserved between humans and mice (**Fig. 1a**), including the region around the AIS-associated SNP (rs169311). We designed three gRNA targeting the 5’ of Xe1, 3’ of Xe1, and 3’ of PEC7 (**Fig. 1a**) and independently injected a pair of gRNA (5’ and 3’ of Xe1, 3’ of Xe1 and 3’ of PEC7, and 5’ of Xe1 and 3’ of PEC7) along with Cas9 protein. We obtained founder lines for all three manipulations, which were further validated by PCR, sequencing, and Southern blot analyses (**Supplementary Fig.2**).

In homozygous Xe1 and Xe1+PEC7 knockout mouse lines, we observed kinks in the distal part of the tail due to a bent caudal vertebra, as observed in micro-CT (**Fig. 2a-b**). This kinky tail phenotype had partial penetrance in Xe1 and Xe1+PEC7 lines (**Table 1a**). Homozygous Xe1 mice had around 20% penetrance of kinky tails in both females and males. Interestingly, homozygous Xe1+PEC7 mice displayed higher penetrance with a significant sex difference, with 60% Xe1+PEC7 females and 42% males having kinky tails (**Table 1a**). To test whether these mice have any additional skeletal abnormalities, we carried out micro-CT at both 4 and 6 months of age for all genotypes, including mice with and without a kinky tail. We did not observe any other apparent skeletal abnormalities, other than the kinky tail (**Fig. 2c, Supplementary Fig. 3**). We also checked the kinky tail progression daily after birth (P0-20, **Fig. 2d**) and found that the tail curvature starts from P0 with bending progressing to the distal part of the tail at later stages (>P7). This is consistent with the PEC7 enhancer activity observed in the neonatal stage (**Fig. 1b, Supplementary Fig. 1a**). In summary, our results suggest that deletion of Xe1 leads to a partially penetrant kinky tail phenotype that is intensified both in terms of penetrance and sex difference when the PEC7 region is deleted along with Xe1.

**Fig. 2.**
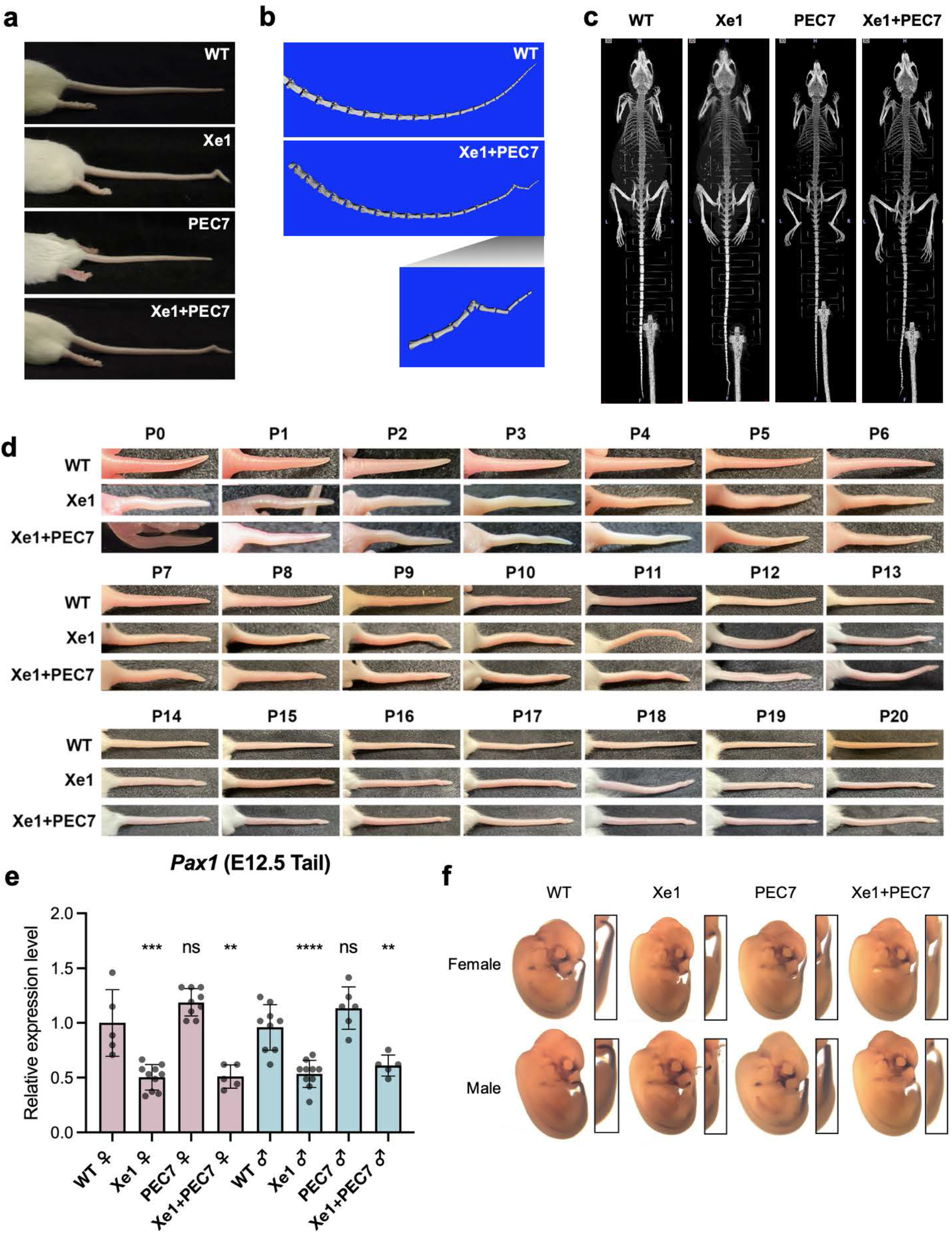
Phenotypic characterization of *Pax1* enhancer knockout mice. **a**, Representative tails for each genotype. **b**, Tail skeletal structure of 2.5-month-old mice scanned by micro-CT (the same mice as shown in **a**). **c**, Whole-body skeletal structure of 6-month-old mice scanned by micro-CT. **d**, Photo of tail from P0 to P20 neonatal stages. **e**, *Pax1* gene expression levels from E12.5 mouse tail as determined by qRT-PCR. Each value represents the ratio of *Pax1* gene expression to that of *β-Actin*, and values are mean ± standard deviation. The expression value of wild type (WT) females was arbitrarily set at 1.0. Each dot represents one embryo and statistical differences were determined using unpaired t test (****<0.001, ***<0.005, **<0.01, ns, not significant). **f**, Whole-mount *in situ* hybridization for *Pax1* of E12.5 mouse embryos.

**Table 1.** Kinky tail ratios for the various mouse genotypes. **a**, Kinky tail ratio observed at 3 weeks of age. Significance was calculated by Fisher’s test (sample number < 5) or Chi-square test (sample number > 5). **b**, Kinky tail ratio observed at 3 weeks of age. Pups were obtained from tamoxifen injected Xe1+PEC7 knockout dams. Significance was calculated by Chi-square test (sample number > 5).

|  |  | Female |  |  |  | Male |  |  |  | P-val. |
| --- | --- | --- | --- | --- | --- | --- | --- | --- | --- | --- |
| Genotype |  | Normal | Kinky | Total | Kinky % | Normal | Kinky | Total | Kinky % | F vs M |
| Xe1 | WT/WT | 31 | 0 | 31 | 0% | 29 | 0 | 29 | 0% | N.S. |
|  | WT/Del | 75 | 2 | 77 | 3% | 75 | 5 | 80 | 6% | N.S. |
|  | Del/Del | 59 | 18 | 77 | 23% | 50 | 11 | 61 | 18% | 0.44 |
| PEC7 | WT/WT | 17 | 0 | 17 | 0% | 21 | 0 | 21 | 0% | N.S. |
|  | WT/Del | 45 | 0 | 45 | 0% | 43 | 0 | 43 | 0% | N.S. |
|  | Del/Del | 30 | 0 | 30 | 0% | 30 | 0 | 30 | 0% | N.S. |
| Xe1+PEC7 | WT/WT | 21 | 0 | 21 | 0% | 30 | 0 | 30 | 0% | N.S. |
|  | WT/Del | 99 | 1 | 100 | 1% | 99 | 2 | 101 | 2% | N.S. |
|  | Del/Del | 23 | 34 | 57 | 60% | 43 | 31 | 74 | 42% | 0.044 |

**Table 1| Kinky tail ratios for the various mouse genotypes.**
|  |  | Female |  |  |  | Male |  |  |  | P-val. |
| --- | --- | --- | --- | --- | --- | --- | --- | --- | --- | --- |
| Genotype |  | Normal | Kinky | Total | Kinky % | Normal | Kinky | Total | Kinky % | F vs M |
| **Xe1+PEC7 del**<br>**Tamoxifen inj.** | Del/Del | 24 | 8 | 32 | 25% | 27 | 10 | 37 | 27% | 0.8484 |

### Xe1 and Xe1 + PEC7 homozygous mice have reduced *Pax1* embryonic tail expression

We next set out to test the expression of *Pax1* in all three lines. As *Pax1* has been reported to be highly expressed at E10.5 to E13.5 in the tail^16,17^, we first analyzed its expression in this tissue. We carried out qRT-PCR and whole-mount *in situ* hybridization (WISH) on E12.5 embryos. We observed, via qRT-PCR, a ∼50% downregulation of *Pax1* expression in Xe1 and Xe1+PEC7 homozygous mice but no significant changes in expression for the PEC7 homozygous mice compared to wild type (WT) mice (**Fig. 2e**). Consistent with the qRT-PCR data, WISH also showed reduction of *Pax1* gene expression in the tail for both these homozygous lines (**Fig. 2f**). We also checked *Pax1* gene expression in ten-week-old thymus and IVD, as it has been reported that *Pax1* is highly expressed in these tissues^26^. PEC7 homozygous female and Xe1 male showed slightly higher expression compared to WT mice in thymus, however other lines did not show a significant *Pax1* gene expression differences in these tissues (**Supplementary Fig. 4**).

### RNA-seq identifies ECM genes to be downregulated in enhancer knockout mice

To identify downstream gene expression changes in the various enhancer knockouts, we performed RNA-seq on E12.5 embryo tails for all three homozygous knockouts (Xe1, PEC7, Xe1+PEC7) and WT mice, each in three females and three males, for a total of 24 samples. We chose this time point, as *Pax1* is strongly expressed at the tail at this time point^16,17.^ We first identified differentially expressed genes between the transcriptomes of male and female mice using the Gene Ontology (GO) term enrichment analysis that is part of the Partek genomics suite (see **Methods**). We found a strong enrichment for ‘embryonic skeletal system morphology’, ‘muscle tissue development’, ‘cell adhesion’, and ‘collagen fibril organization’ in females (**Fig. 3a**). Xe1+PEC7 females showed the highest association with ‘embryonic skeletal system morphologies’.

**Fig. 3.**
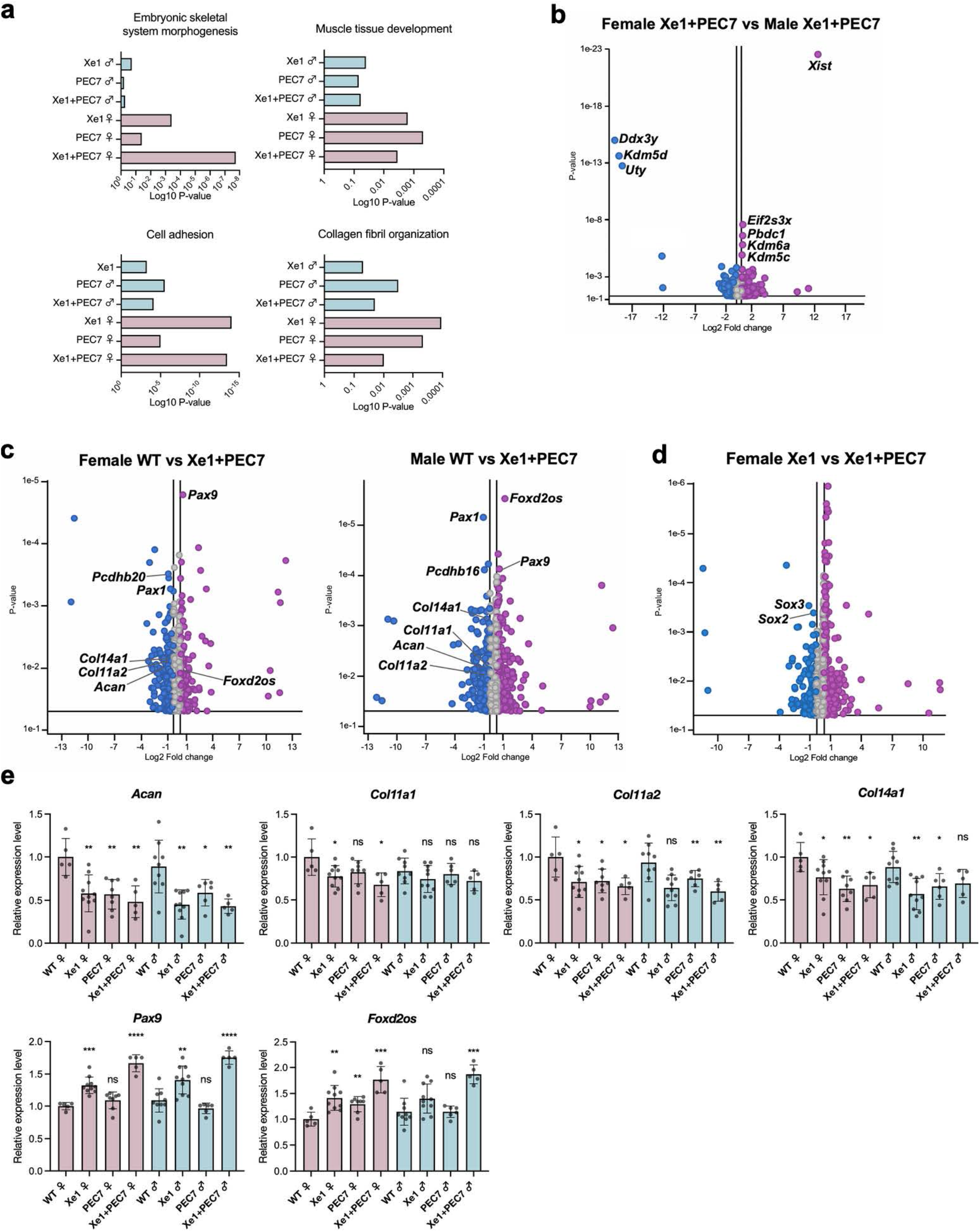
Gene expression profiling of E12.5 tails by RNA-seq and qPCR. **a**, Enriched GO terms for gene sets expressed differentially between wild type (WT) and enhancer knockout mice (*P*≦0.05). **b-d**, Volcano plots showing the global transcriptional changes for the indicated groups. Each circle represents one gene. The log2 fold change in the indicated genotype is represented on the X-axis. The Y-axis shows the p-value. A p-value of 0.05 and a fold change of 1.5 are indicated by lines. **e**, Gene expression levels dissected from E12.5 mouse tail as determined by qRT-PCR. Each value represents the ratio of gene expression to that of *β-Actin* and values are mean ± standard deviation. The expression value of WT females was arbitrarily set at 1.0. Each dot represents one embryo and statistical differences were determined using unpaired t test (****<0.001, ***<0.005, **<0.01, ns, not significant).

We next set out to identify differentially expressed genes between males and females, specifically in the Xe1+PEC7 homozygous knockouts to identify candidate genes that might be responsible for the sex differences observed for the kinky tail penetrance. Analysis of differentially expressed genes between males and females identified only sex-chromosome differential genes (**Fig. 3b**). Next, we compared the gene expression differences between WT and Xe1+PEC7 in both sexes separately.

We observed significant downregulation of *Pax1* in the Xe1+PEC7 knockouts as expected (**Fig. 3c**). In addition, we saw downregulation of extracellular matrix genes (*Acan, Col11a1, Col11a2* and *Col14a1*) and protocadherin genes involved in cell-adhesion (*Pcdhb16* and *Pcdhb20*). Conversely, we observed upregulation of *Pax9* and *Foxd2os* in both sexes. *Pax9* and *Pax1* has redundant function in axial skeletogenesis^16,18,27^ and *Pax9* expression was shown to be upregulated to compensate for *Pax1* expression in *Pax1* knockout mice^27^ fitting with our observation of lower *Pax1* expression in our qRT-PCR and RNA-seq. *FOXD2-AS1* (human ortholog of *Foxd2os*) is known to be highly expressed and regulates chondrocyte proliferation in osteoarthritis patients^28,29^. We next compared the gene expression changes between Xe1 and Xe1+PEC7 homozygous female mice, as Xe1 homozygous mice did not have a kinky tail sex bias phenotype. We found *Sox2* and *Sox3* to be downregulated in Xe1+PEC7 knockout females compared to Xe1 knockout females (**Fig. 3d**). *Sox2* and *Sox3* are known to have a redundant function in the development of otic and epibranchial tissue^30^ and *Sox2* is involved in mesoderm differentiation in zebrafish tail buds^31^.

To validate our RNA-seq results and analyze specific genotypes in more detail, we performed qRT-PCR using at least five mice for all genotypes and sex. We first confirmed that the extracellular matrix genes (*Acan, Col11a1, Col11a2*, and *Col14a1*) are downregulated in all homozygous knockout mouse lines (**Fig. 3e**). Interestingly, in the PEC7 homozygous knockouts we also observed downregulation of *Acan, Col11a2*, and *Col14a1* (**Fig. 3e**) despite not observing any expression changes in *Pax1* in these mice (**Fig. 2e**). *Pax9* and *Foxd2os* were both upregulated in Xe1 and Xe1+PEC7 homozygous knockout mice. Interestingly, upregulation of both these genes was higher in Xe1+PEC7 mice compared to Xe1 (**Fig. 3e**), although *Pax1* expression was similar in these mice (**Fig. 2e**). In PEC7 homozygous mice, we only observed *Foxd2os* to be significantly upregulated in females. Combined, these results suggest that Xe1 and PEC7 might work synergistically and that the deletion of PEC7 might affect the expression of other genes that are not directly regulated by *Pax1*.

### Tamoxifen injection in Xe1+PEC7 pregnant mothers abolishes kinky tail sex difference

As this region is associated with AIS only in females^10^ and female Xe1+PEC7 knockout mice show higher kinky tail penetrance, we hypothesized that estrogen could be involved in this process, based on previous reports associating it with AIS^32^. Previous work showed that tamoxifen, a selective estrogen receptor modulator, decreases the rate of curve progression in a melatonin deficient bipedal scoliosis mouse model^33^. We thus injected tamoxifen into Xe1+PEC7 knockout E17.5 pregnant females and compared their kinky tail phenotypic ratio of pups at day 21 to uninjected mice. Interestingly, we observed significantly reduced penetrance for the kinky tail phenotype and that the previous sex difference for Xe1+PEC7 knockout was ablated, showing an almost equal number of males and females with a kinky tail, 25% in females and 27% in males (**Table 1b**) compared to 60% and 42% in non-injected mice (**Table 1a**).

To further characterize the genes involved in this process, we conducted RNA-seq using the distal part of the P2 tail tip, due to it being the timepoint and location where the kinky tail initiates (**Fig. 2d**) and also the tamoxifen injected condition. We extracted RNA from WT, Xe1, PEC7, Xe1+PEC7 and Xe1+PEC7 pups from tamoxifen injected dams (Xe1+PEC7(Tam)). We used four samples per condition and sex, totaling 40 samples for RNA-seq. In contrast to our E12.5 analyses, we observed no significant gene expression changes for *Pax1* between the different genotypes and sexes when compared to WT at this P2 time point (**Fig. 4a**). These results suggest that the functional effect and activity of Xe1 and/or PEC7 is likely earlier. Next, we compared males and females to identify differential GO term enrichment. We found Xe1+PEC7 specific enrichment in females for ‘Dynein heavy chain binding’ and ‘Ciliary based body organization’ (**Fig. 4b**). Intriguingly, knockout of axonemal dynein and its assembly factor genes, such as *dnah10, dnaaf1* and *zmynd10*, are known to lead to a scoliosis phenotype in zebrafish^34,35^. Defects of motile cilia were also shown to cause scoliosis in zebrafish^36^. Next, to identify differentially expressed genes in these processes, we compared gene expression differences between males and females in Xe1+PEC7 knockout mice, finding only sex chromosome associated genes (**Fig. 4c**). Then, we focused only on females and analyzed gene expression differences between the various genotypes, including WT versus Xe1+PEC7, and Xe1 versus Xe1+PEC7. Interestingly, *Foxj1* and *Efcab1*, that are involved with cilium function^37,38,39^, were found to be upregulated in Xe1+PEC7 female mice compared to WT females (**Fig. 4d**). We also found *Dnaaf1* to be downregulated in Xe1+PEC7 (**Fig. 4d-e**). The dynein axonemal heavy chain (DNAH) family genes, such as *Dnah11* and *Dnah3*, were differentially expressed (**Fig. 4d-e**). In addition, *Mmp9*, an enzyme involved in cartilage degradation during endochondral ossification^40^, was found to be downregulated in Xe1+PEC7 female mice compared to Xe1 female mice.

**Fig. 4.**
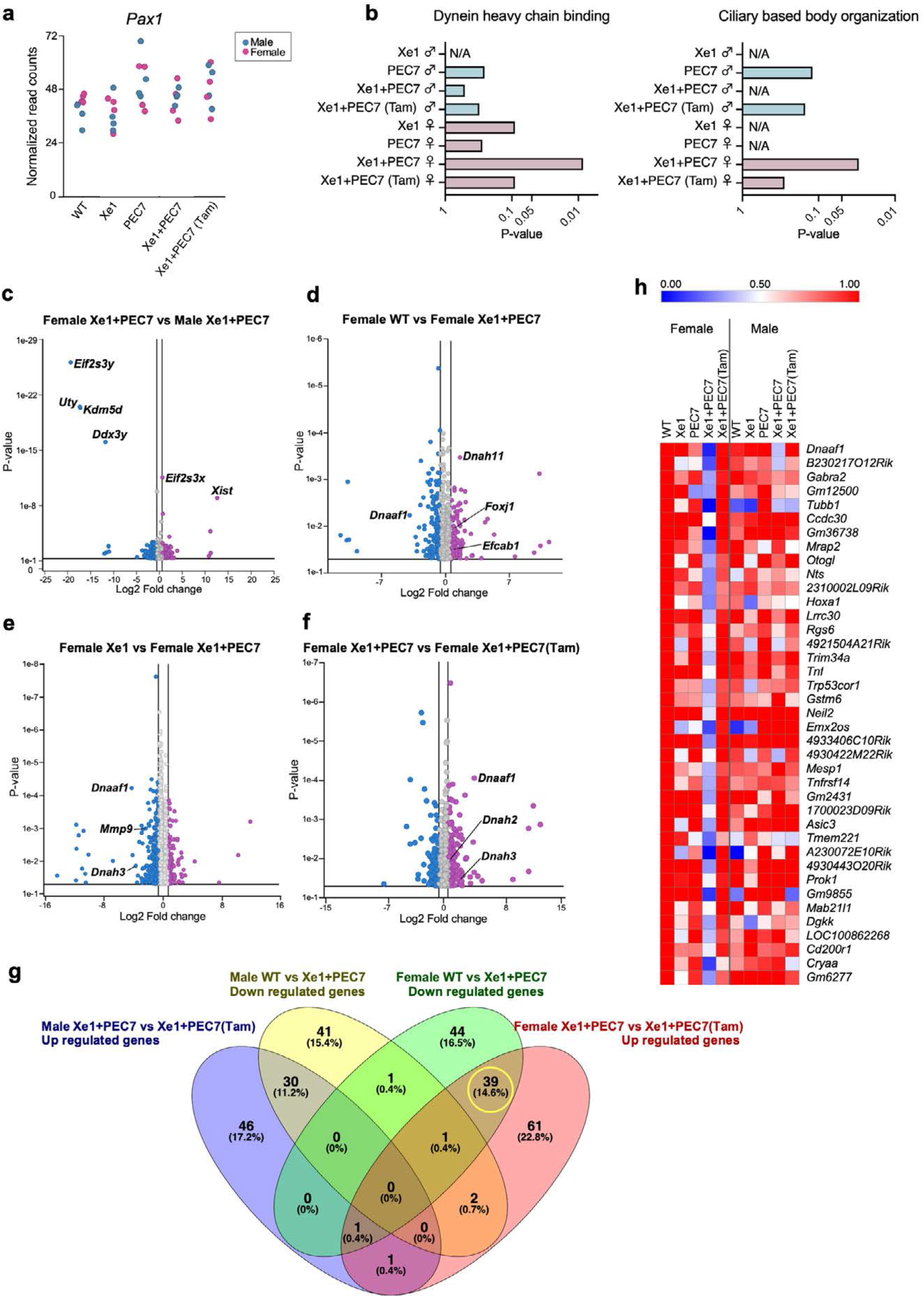
Gene expression profiling at P2 tail by RNA-seq. **a**, *Pax1* gene expression levels from RNA-seq data. **b**, Enriched GO terms for gene sets differentially expressed between wild type (WT) and enhancer knockout mice (*P*≦0.05). **c-f**, Volcano plots showing the global transcriptional changes for the various groups. Each circle represents one gene. The log2 fold change in the indicated genotype is represented on the X-axis. The Y-axis shows the *P*-value. A p-value of 0.05 and a fold change of 1.5 are indicated by lines. **g**, Venn diagram showing genes uses the following criteria: downregulated genes compared between WT and Xe1+PEC7 male or female and upregulated genes compared between Xe1+PEC7 and Xe1+PEC7(Tam) male or female (at least ± 2-fold changes; *P*≦0.05). Each number shows gene number and percentage (%). **h**, heat map of 39 differentially expressed genes identified from the Venn diagram. The expression value of the WT female group was arbitrarily set at 1.0.

We next set out to assess the effects of tamoxifen on gene expression. We first compared Xe1+PEC7 female mice with or without tamoxifen. Intriguingly, *Dnaaf1, Dnah2*, and *Dnah3* expression levels were upregulated in the Xe1+PEC7(Tam) females (**Fig. 4f**). These results suggest that dynein and cilia function could be associated with kinky tail progression in Xe1+PEC7 knockout females. To further identify genes that could be involved in the sex difference phenotype in Xe+PEC7 mice, we characterized genes that are downregulated in Xe1+PEC7 knockout mice compared to WT, or upregulated in Xe1+PEC7(Tam) mice compared to uninjected mice. We identified 39 genes that were downregulated in Xe1+PEC7 knockout and restored with tamoxifen injection specifically in females (**Fig. 4g-h**). While we did not observe any specific GO enriched term for these genes, we did identify some interesting AIS-associated genes. They include the aforementioned *Dnaaf1*, the deletion of which causes a scoliosis phenotype in zebrafish^34^; *Hoxa1*, a homeobox gene essential for the development of head and neck structures, including hindbrain, ear, and occipital and hyoid bones^41,42^; *Mesp1*, a transcription factor involved in mesoderm specification, somite boundary formation and somite polarity regulation^43,44^; and *Tnfrsf14*, a receptor for *Tnfsf14. Tnfsf14* is primarily expressed in lymphocytes, as a critical regulator of key enzymes that control lipid metabolism^45^, and it resides in a region on chromosome 19p13.3 that has been linked to AIS^46^. In summary, our RNA-seq identified genes that show differential expression between the various genotypes and with or without tamoxifen treatment, several of which are involved in ciliary or dynein function and could be involved in AIS sexual dimorphic phenotypes.

## Discussion

Using mouse transgenics and knockouts we characterized an AIS susceptibility locus downstream of *PAX1* that is associated with female AIS^10^. We found that while both Xe1 and PEC7 function as enhancers, only removal of Xe1 leads to a kinky tail phenotype. Interestingly though, removal of PEC7 along with Xe1 leads both to an increased penetrance of this phenotype and sexual dimorphism, with females showing a higher prevalence. Using RNA-seq, we extensively characterized the genes and pathways associated with this phenotype, finding genes involved in ECM, ciliary or dynein function. Using tamoxifen injections in Xe1+PEC7 pregnant females, we show that the sex bias is likely associated with estrogen signaling. Combined, our results further dissect this female AIS-associated locus and identify candidate genes and pathways for AIS.

Modeling AIS in mice had been limited^47^. This is likely due to mice being quadrupedal compared to bipedal humans, leading to inherent differences of their spinal anatomy. A kinky tail phenotype has been observed for several genes associated with scoliosis^47^, including *Pax1*^19^. While *Pax1* homozygous mice also show lumbar scoliosis, we only observed a kinky tail in our enhancer knockout mice. In addition, the kinky tail in *Pax1* homozygous knockout phenotype shows complete penetrance and is much more severe, with several bones affected throughout the tail^19^. Our enhancer knockout leads to a less severe phenotype with reduced penetrance, in line with previous enhancer knockout phenotypes which usually tend to show a more subtle condition than that of their target gene knockout^48^. This could be due to genes having multiple functions in various tissues versus an enhancer being tissue/cell-type specific^49^, promoters having a stronger regulatory role than enhancers and/or enhancer redundancy^50,51^, i.e. having multiple enhancers for a specific target gene with similar function. This more subtle and not fully penetrant phenotype is also in line with AIS-associated GWAS variants being associated with AIS susceptibility, where having a specific variant along with environmental and/or other genetic factors combined lead to AIS.

Our mouse enhancer assays showed that PEC7 functions as an enhancer in the distal part of the neonate tail and adult IVD. However, its removal did not lead to an observable phenotype, as assessed by microCT up to six months of age. In contrast, independent removal of Xe1 led to a kinky tail phenotype with 20% penetrance with no differences between males and females. In line with this locus being associated with AIS in females^10^, we observed an increased penetrance of a kinky tail phenotype in females only in our Xe1+PEC7 homozygous mice. While Xe1 does not contain an AIS-associated variant, the homologous human PEC7 region does, suggesting that PEC7 could be the cause of the sexual dimorphism in this region. Interestingly, this locus has also been associated with male pattern baldness^52,53^, suggesting that it can also affect male specific phenotypes. Our tamoxifen injections into E17.5 pregnant Xe1+PEC7 significantly reduced the kinky tail penetrance in both males and even more in females, bringing it to similar levels as Xe1 knockouts, suggesting that PEC7 might be regulated by estrogen signaling. Utilizing JASPAR^54^, we find that the mouse PEC7 region contains an estrogen receptor 2 (*ESR2*) motif (**Fig. 1a**). However, this motif is not conserved in humans nor did we find other ESR2 motifs in the human PEC7 sequence. Further characterization of how this region leads to various sexual dimorphic phenotypes, including scoliosis or male pattern baldness, would be of extreme interest.

Our RNA-seq analyses in both E12.5 and P2 tails identified numerous genes and pathways that could be linked to AIS in general and to a sexual dimorphic phenotype. We observed several ECM genes (*Acan, Col11a1, Col11a2, Col14a1, Pcdhb16* and *Pcdhb20*) to be significantly downregulated in both Xe1+PEC7 male and female knockouts. The ECM has been strongly associated with AIS, with many other AIS GWAS loci residing near ECM-associated genes and zebrafish ECM gene knockouts having a scoliosis phenotype^55^. In terms of sexually dimorphic genes, we analyzed both Xe1 versus Xe1+PEC7 homozygous female mice, as Xe1 homozygous did not have a kinky tail sex bias, and tamoxifen treated versus untreated Xe1+PEC7 knockout P2 tails, as tamoxifen treatment demolished the kinky tail sex bias in these Xe1+PEC7 knockouts. In particular, we identified the dynein-associated gene, *Dnaaf1*, which is known to cause scoliosis in zebrafish^34^, to be downregulated in Xe1+PEC7 compared to Xe1 female knockout P2 tails. In addition, defects of motile cilia are known to cause scoliosis in zebrafish^36^, fitting with our observation of several cilia-associated genes. We also found *Sox2* and *Sox3* which are known to be involved in the development of otic and epibranchial tissue^30^ and mesoderm differentiation in zebrafish tail buds^31^, to be downregulated in Xe1+PEC7 homozygous female mice compared to Xe1 homozygous female mice at E12.5 tails. In addition, for *SOX3*, altered gene regulation, due to an interchromosomal insertion downstream of this gene, is thought to lead to congenital generalized hypertrichosis along with spina bifida and scoliosis^56^. As their name implies, sex determining region Y (SRY)-box transcription factors have been associated with various sex dimorphic phenotypes. *SOX2* for example is known to be involved in sexual dimorphic differences in the peripheral nervous system^57^, which has been shown to have a major role in scoliosis etiology^58^. Interestingly, *Tnfrsf14*, which was found to be significantly downregulated in Xe1+PEC7 homozygous knockouts but not in the tamoxifen treatment, was found to be associated with sexual dimorphic expression in osteoprogenitors in progesterone receptor knockout mice^59^.

Gene regulatory elements were found to be a major cause of sex associated gene expression^60^, and sexual dimorphic phenotypes, such as inguinal hernia^61^ or cancer^62^. Further work dissecting the sexual dimorphic function and mechanism of these genes and regulatory elements could shed light on AIS prevalence differences between males and females specifically and in general for other sexual dimorphic phenotypes.

## Methods

### Generation of transgenic mice

For the mouse transgenic enhancer assay, we used an *Hsp68*-mCherry (hCR) plasmid^63^, a kind gift provided by Drs. Len Pennacchio, Axel Visel and Dianne Dickel (LBNL). This plasmid was digested with *Kpn*I and the PCR fragment was cloned, including multi cloning sites amplified by HSP68-MCS primer set (**Supplementary Table 1**), using *Hsp68*-LacZ plasmid as a template^25^. This modified Hsp68-mCherry plasmid was digested with *Hind*III, and mouse PEC7 (chr2:147,566,137-147,567,924; *mm10*) amplified by PEC7-HSP68 primer set with FVB/NJ mouse genome as a template was cloned into the plasmid. The plasmid was digested with *Sal*I, and the assayed DNA fragment was released from the backbone vector and used for pronuclear injection, which was performed by the Transgenic Gene Targeting Core at the Gladstone Institute. All mouse work was approved by the UCSF IACUC protocol number AN197608 and was conducted in accordance with AALAC and NIH guidelines. FVB/NJ mouse stain (Jackson Laboratory; 001800) was used.

### Generation and validation of knockout mice

Mouse Xe1 (chr2:147,560,271-147,561,812; *mm10*) and PEC7 (chr2:147,566,137-147,567,924; *mm10*) homology sequence was identified with Ensemble genome browser^24^. To generate Xe1, PEC7 or Xe1+PEC7 knockout mice, three gRNAs were designed (**Fig.1a, Supplementary Table 1**) using the gRNA design tool on the Integrated DNA Technologies (IDT) website and selected based on low off-target and high on-target scores. Two crRNA, tracrRNA and Cas9 protein (IDT; 1081058) were injected to zygotes via the Transgenic Gene Targeting Core at the Gladstone Institute. All mouse work was approved by the UCSF IACUC, protocol number AN197608, and was conducted in accordance with AALAC and NIH guidelines. FVB/NJ mouse stain (Jackson Laboratory, catalog no. 001800) was used.

PCR-Sanger sequencing (primers provided in **Supplementary Table 1**) was performed using standard techniques. For Southern blot analyses, genomic DNA was treated with *Bgl*Il or *Pvu*II (New England Biolabs; R0144 or R0151) and fractionated by agarose gel electrophoreses. Following capillary transfer onto nylon membranes, blots were hybridized with Digoxigenin (DIG)-labeled DNA probes (**Supplementary Table 2**) amplified by the PCR DIG Probe Synthesis Kit (Sigma-Aldrich; 11636090910). The hybridized probe was immunodetected with anti-digoxigenin Fab fragments conjugated to alkaline phosphatase (Sigma-Aldrich; 11093274910) and visualized with a CDP star (Sigma-Aldrich; 11685627001), according to the manufacturer’s protocol. Chemiluminescence was detected using FluorChem E (ProteinSimple; 92-14860-00).

### Micro-computed tomography (microCT)

MicroCT scans were performed on fixed mouse skeletons using a Scanco Medical μCT50 at the UCSF Core Center of Musculoskeletal Biology and Medicine. Specimens were scanned at 30.0 micron resolution with scanner settings of 55kVP, 109μA, 6W, 0.5mm AI filter as well as an integration time of 500ms. Reconstructions were converted into DICOM files using Scanco Medical’s integrated μCT Evaluation Program V6.5-3, then converted into 3D volumes using μCT Ray V4.0-4. MicroCT scans were performed on living mice with U-CT (MILabs B.V.), part of VECT or 4CT at the pre-Clinical Imaging core in the Department of Radiology and Biomedical Imaging at UCSF. The scanning parameters were: x-ray tube voltage at 55 kVp and current at 0.19 mA and 75 ms exposure time at each angular step over 360 degrees and 480 steps. Once the data were acquired, the manufacturer-provided cone beam FDK algorithm was used for image reconstruction with the voxel size of 0.160 mm x 0.160 mm x 0.160 mm.

### Quantitative RT-PCR

Total RNA was collected from mouse tissues using TRIzol (Thermo Fisher Scientific; 15596026) and converted to cDNA using ReverTra Ace qPCR-RT master mix with genomic DNA (gDNA) remover (Toyobo; FSQ-301). qPCR was performed using SsoFast EvaGreen supermix (Bio Rad; 1725205). Primer sequences used for qPCR are listed in **Supplementary Table 3**.

### Whole-mount *in situ* hybridization

Mouse E12.5 embryos were fixed in 4% paraformaldehyde. A plasmid containing mouse *Pax1* (GenScript; OMu21524) was used as template for DIG-labeled probes. Mouse whole-mount *in situ* hybridization was performed according to standard procedures^64^. Briefly, embryos were rehydrated and treated with Proteinase K (Promega; V3021). Following re-fixation and prehybridization, embryos were hybridized with a DIG-labeled probe. The hybridized probe was immunodetected with anti-digoxigenin Fab fragments conjugated to alkaline phosphatase (Sigma-Aldrich; 11093274910) and visualized with a Bmpurple (Sigma-Aldrich; 11442074001) according to the manufacturer’s protocol.

### Tamoxifen injection

Tamoxifen (Sigma; T5648) was dissolved in sunflower oil (Sigma; S5007d). Pregnant females were injected with tamoxifen subcutaneously (30 mg/kg body weight) at gestational day 17.5. Pups’ tail morphology was visually analyzed at P21.

### RNA-seq

Total RNA was extracted as described above. RNA-seq was conducted by Novogene. The library was generated with NEBNext Ultra II RNA Library Prep Kit for Illumina (NEB; 7770), and sequencing was done using the NovaSeq 6000 S4 platform with PE150. The data were analyzed using Partek Flow (Version 10.0). After primary quality assessment was performed, bases and reads with low quality were filtered out and the reads were aligned to mouse reference genome (mm10) using the STAR (Version 2.7.8a)^65^. The final BAM files were quantified using the Partek E/M algorithm^66^.

Normalization of read count was performed by the total number of counts (count per million) plus 0.0001, and all genes with less than ten normalized read counts were excluded from subsequent analyses. Differentially expressed genes were identified using Partek gene-specific analysis (GSA) algorithm. Geneotology^67^ and Morpheus (RRID: SCR_017386) were used for data analysis.

## Data availability

All RNA-seq data are available from Sequence Read Archive (SRA), accession number PRJNA951902.

## Acknowledgments

This work was supported in part by the National Institute of Child Health and Development (NICHD) grant number 1P01HD084387 (CAW and NA). AU was supported by the Japan Society for the Promotion of Science (JSPS) postdoctoral fellowships for research abroad, the Uehara Memorial Foundation postdoctoral fellowship and the National Human Genome Research Institute (NHGRI).

## Author contributions

Conceptualization: AU, RRS, NA

Methodology: AU, RRS, YZ, JZ, MN, EM, XR

Samples: AU, RRS, YZ, JZ, MN, EM, XR

Writing, Review & Editing: AU, RRS, JJR, CAW, NA with contributions from all authors

Visualization: AU, RRS, NA

Supervision: NA

Funding acquisition: NA, CAW

## References

1. Weinstein, S. L., Dolan, L. A., Wright, J. G. & Dobbs, M. B. Effects of bracing in adolescents with idiopathic scoliosis. N. Engl. J. Med. 369, 1512–1521 (2013).

2. Raggio, C. L. Sexual dimorphism in adolescent idiopathic scoliosis. Orthop. Clin. North Am. 37, 555–558 (2006).

3. Karol, L. A., Johnston, C. E., 2nd, Browne, R. H. & Madison, M. Progression of the curve in boys who have idiopathic scoliosis. J. Bone Joint Surg. Am. 75, 1804–1810 (1993).

4. Grauers, A., Rahman, I. & Gerdhem, P. Heritability of scoliosis. Eur. Spine J. 21, 1069–1074 (2012).

5. Khanshour, A. M. et al Genome-wide meta-analysis and replication studies in multiple ethnicities identify novel adolescent idiopathic scoliosis susceptibility loci. Hum. Mol. Genet. 27, 3986–3998 (2018).

6. Kou, I. et al Genetic variants in GPR126 are associated with adolescent idiopathic scoliosis. Nat. Genet. 45, 676–679 (2013).

7. Londono, D. et al A meta-analysis identifies adolescent idiopathic scoliosis association with LBX1 locus in multiple ethnic groups. J. Med. Genet. 51, 401–406 (2014).

8. Miyake, A. et al Identification of a susceptibility locus for severe adolescent idiopathic scoliosis on chromosome 17q24.3. PLoS One 8, e72802 (2013).

9. Ogura, Y. et al A functional SNP in BNC2 is associated with adolescent idiopathic scoliosis. Am. J. Hum. Genet. 97, 337–342 (2015).

10. Sharma, S. et al A PAX1 enhancer locus is associated with susceptibility to idiopathic scoliosis in females. Nat. Commun. 6:6452.,10.1038/ncomms7452. (2015).

11. Sharma, S. et al Genome-wide association studies of adolescent idiopathic scoliosis suggest candidate susceptibility genes. Hum. Mol. Genet. 20, 1456–1466 (2011).

12. Takahashi, Y. et al A genome-wide association study identifies common variants near LBX1 associated with adolescent idiopathic scoliosis. Nat. Genet. 43, 1237–1240 (2011).

13. Zhu, Z. et al Genome-wide association study identifies new susceptibility loci for adolescent idiopathic scoliosis in Chinese girls. Nat. Commun. 6, 8355 (2015).

14. Kou, I. et al Genome-wide association study identifies 14 previously unreported susceptibility loci for adolescent idiopathic scoliosis in Japanese. Nat. Commun. 10, 3685 (2019).

15. Makki, N. et al Genomic characterization of the adolescent idiopathic scoliosis-associated transcriptome and regulome. Hum. Mol. Genet. 29, 3606–3615 (2021).

16. Wallin, J. et al The role of Pax-1 in axial skeleton development. Development 120, 1109–1121 (1994).

17. Sivakamasundari, V., Kraus, P., Jie, S. & Lufkin, T. Pax1(EGFP): new wildtype and mutant EGFP mouse lines for molecular and fate mapping studies. Genesis 51, 420–429 (2013).

18. Wilm, B., Dahl, E., Peters, H., Balling, R. & Imai, K. Targeted disruption of Pax1 defines its null phenotype and proves haploinsufficiency. Proc. Natl. Acad. Sci. U. S. A. 95, 8692–8697 (1998).

19. Adham, I. M. et al The scoliosis (sco) mouse: a new allele of Pax1. Cytogenet. Genome Res. 111, 16–26 (2005).

20. Sivakamasundari, V. et al A developmental transcriptomic analysis of Pax1 and Pax9 in embryonic intervertebral disc development. Biol. Open 6, 187–199 (2017).

21. Takimoto, A. et al Differential transactivation of the upstream aggrecan enhancer regulated by PAX1/9 depends on SOX9-driven transactivation. Sci. Rep. 9, 4605 (2019).

22. Kokubu, C. et al A transposon-based chromosomal engineering method to survey a large cisregulatory landscape in mice. Nat. Genet. 41, 946–952 (2009).

23. ENCODE Project Consortium. An integrated encyclopedia of DNA elements in the human genome. Nature 489, 57–74 (2012).

24. Cunningham, F. et al Ensembl 2022. Nucleic Acids Res. 50, D988–D995 (2022).

25. Kothary, R. et al A transgene containing lacZ inserted into the dystonia locus is expressed in neural tube. Nature 335, 435–437 (1988).

26. Wallin, J. et al Pax1 is expressed during development of the thymus epithelium and is required for normal T-cell maturation. Development 122, 23–30 (1996).

27. Peters, H. et al Pax1 and Pax9 synergistically regulate vertebral column development. Development 126, 5399–5408 (1999).

28. Wang, Y., Cao, L., Wang, Q., Huang, J. & Xu, S. LncRNA FOXD2-AS1 induces chondrocyte proliferation through sponging miR-27a-3p in osteoarthritis. Artif. Cells Nanomed. Biotechnol. 47, 1241–1247 (2019).

29. Cao, L., Wang, Y., Wang, Q. & Huang, J. LncRNA FOXD2-AS1 regulates chondrocyte proliferation in osteoarthritis by acting as a sponge of miR-206 to modulate CCND1 expression. Biomed. Pharmacother. 106, 1220–1226 (2018).

30. Gou, Y., Guo, J., Maulding, K. & Riley, B. B. Sox2 and sox3 cooperate to regulate otic/epibranchial placode induction in zebrafish. Dev. Biol. 435, 84–95 (2018).

31. Kinney, B. A. et al Sox2 and canonical Wnt signaling interact to activate a developmental checkpoint coordinating morphogenesis with mesoderm fate acquisition. Cell Rep. 33, 108311 (2020).

32. Zheng, S. et al Estrogen promotes the onset and development of idiopathic scoliosis via disproportionate endochondral ossification of the anterior and posterior column in a bipedal rat model. Exp. Mol. Med. 50, 1–11 (2018).

33. Demirkiran, G. et al Selective estrogen receptor modulation prevents scoliotic curve progression: radiologic and histomorphometric study on a bipedal C57Bl6 mice model. Eur. Spine J. 23, 455– 462 (2014).

34. Wang, Y., Troutwine, B. R., Zhang, H. & Gray, R. S. The axonemal dynein heavy chain 10 gene is essential for monocilia motility and spine alignment in zebrafish. Dev. Biol. 482, 82–90 (2022).

35. Wang, Y. et al Coding variants coupled with rapid modeling in zebrafish implicate dynein genes, dnaaf1 and zmynd10, as adolescent idiopathic scoliosis candidate genes. Front. Cell Dev. Biol. 8, 582255 (2020).

36. Grimes, D. T. et al Zebrafish models of idiopathic scoliosis link cerebrospinal fluid flow defects to spine curvature. Science 352, 1341–4. doi: 10.1126/science.aaf6419. (2016).

37. Yu, X., Ng, C. P., Habacher, H. & Roy, S. Foxj1 transcription factors are master regulators of the motile ciliogenic program. Nat. Genet. 40, 1445–1453 (2008).

38. Stubbs, J. L., Oishi, I., Izpisúa Belmonte, J. C. & Kintner, C. The forkhead protein Foxj1 specifies node-like cilia in Xenopus and zebrafish embryos. Nat. Genet. 40, 1454–1460 (2008).

39. Sasaki, K. et al Calaxin is required for cilia-driven determination of vertebrate laterality. Commun. Biol. 2, 226 (2019).

40. Kojima, T. et al Histochemical aspects of the vascular invasion at the erosion zone of the epiphyseal cartilage in MMP-9-deficient mice. Biomed. Res. 34, 119–128 (2013).

41. Tischfield, M. A. et al Homozygous HOXA1 mutations disrupt human brainstem, inner ear, cardiovascular and cognitive development. Nat. Genet. 37, 1035–1037 (2005).

42. Muscarella, L. A. et al HOXA1 gene variants influence head growth rates in humans. Am. J. Med. Genet. B Neuropsychiatr. Genet. 144B, 388–390 (2007).

43. Ajima, R., Sakakibara, Y., Sakurai-Yamatani, N., Muraoka, M. & Saga, Y. Formal proof of the requirement of MESP1 and MESP2 in mesoderm specification and their transcriptional control via specific enhancers in mice. Development 148, (2021).

44. Takahashi, Y. et al Differential contributions of Mesp1 and Mesp2 to the epithelialization and rostro-caudal patterning of somites. Development 132, 787–796 (2005).

45. Lo, J. C. et al Lymphotoxin ß Receptor□Dependent control of lipid homeostasis. Science 316, 285–288 (2007).

46. Chan, V. et al A genetic locus for adolescent idiopathic scoliosis linked to chromosome 19p13.3. Am. J. Hum. Genet. 71, 401–406 (2002).

47. Ouellet, J. & Odent, T. Animal models for scoliosis research: state of the art, current concepts and future perspective applications. Eur. Spine J. 22 Suppl 2, S81–95 (2013).

48. Chatterjee, S. & Ahituv, N. Gene Regulatory Elements, Major Drivers of Human Disease. Annu. Rev. Genomics Hum. Genet. 18:45–63., 10.1146/annurev-genom-091416-035537. Epub 2017 Apr 7. (2017).

49. Heintzman, N. D. et al Histone modifications at human enhancers reflect global cell-type-specific gene expression. Nature 459, 108–112 (2009).

50. Osterwalder, M. et al Enhancer redundancy provides phenotypic robustness in mammalian development. Nature 554, 239–243 (2018).

51. Hong, J. W., Hendrix, D. A. & Levine, M. S. Shadow enhancers as a source of evolutionary novelty. Science 321, 1314. (2008).

52. Richards, J. B. et al Male-pattern baldness susceptibility locus at 20p11. Nat. Genet. 40, 1282–1284 (2008).

53. Hillmer, A. M. et al Susceptibility variants for male-pattern baldness on chromosome 20p11. Nat. Genet. 40, 1279–1281 (2008).

54. Fornes, O. et al JASPAR 2020: update of the open-access database of transcription factor binding profiles. Nucleic Acids Res. 48, D87–D92 (2020).

55. Wise, C. A. et al The cartilage matrisome in adolescent idiopathic scoliosis. Bone Res. 8, 13 (2020).

56. Zhu, H. et al X-linked congenital hypertrichosis syndrome is associated with interchromosomal insertions mediated by a human-specific palindrome near SOX3. Am. J. Hum. Genet. 88, 819– 826 (2011).

57. Chernov, A. V. & Shubayev, V. I. Sexually dimorphic transcriptional programs of early-phase response in regenerating peripheral nerves. Front. Mol. Neurosci. 15, 958568 (2022).

58. Blecher, R. et al The proprioceptive system masterminds spinal alignment: Insight into the mechanism of scoliosis. Dev. Cell 42, 388–399.e3 (2017).

59. Kot, A. et al Sex dimorphic regulation of osteoprogenitor progesterone in bone stromal cells. J. Mol. Endocrinol. 59, 351–363 (2017).

60. Oliva, M. et al The impact of sex on gene expression across human tissues. Science 369, (2020).

61. Choquet, H. et al Ancestry- and sex-specific effects underlying inguinal hernia susceptibility identified in a multiethnic genome-wide association study meta-analysis. Hum. Mol. Genet. 31, 2279–2293 (2022).

62. Kfoury, N. et al Brd4-bound enhancers drive cell-intrinsic sex differences in glioblastoma. Proc. Natl. Acad. Sci. U. S. A. 118, e2017148118 (2021).

63. Dickel, D. E. et al Ultraconserved enhancers are required for normal development. Cell 172, 491–499.e15 (2018).

64. Hargrave, M., Bowles, J. & Koopman, P. In situ hybridization of whole-mount embryos. Methods Mol. Biol. 326, 103–113 (2006).

65. Dobin, A. et al STAR: ultrafast universal RNA-seq aligner. Bioinformatics 29, 15–21 (2013).

66. Xing, Y. et al An expectation-maximization algorithm for probabilistic reconstructions of fulllength isoforms from splice graphs. Nucleic Acids Res. 34, 3150–3160 (2006).

67. Ashburner, M. et al Gene ontology: tool for the unification of biology. The Gene Ontology Consortium. Nat. Genet. 25, 25–29 (2000).

