## Supplementary material for "Deletion of *Pax1* scoliosis-associated regulatory elements leads to a female-biased tail abnormality": Supplmentary materials

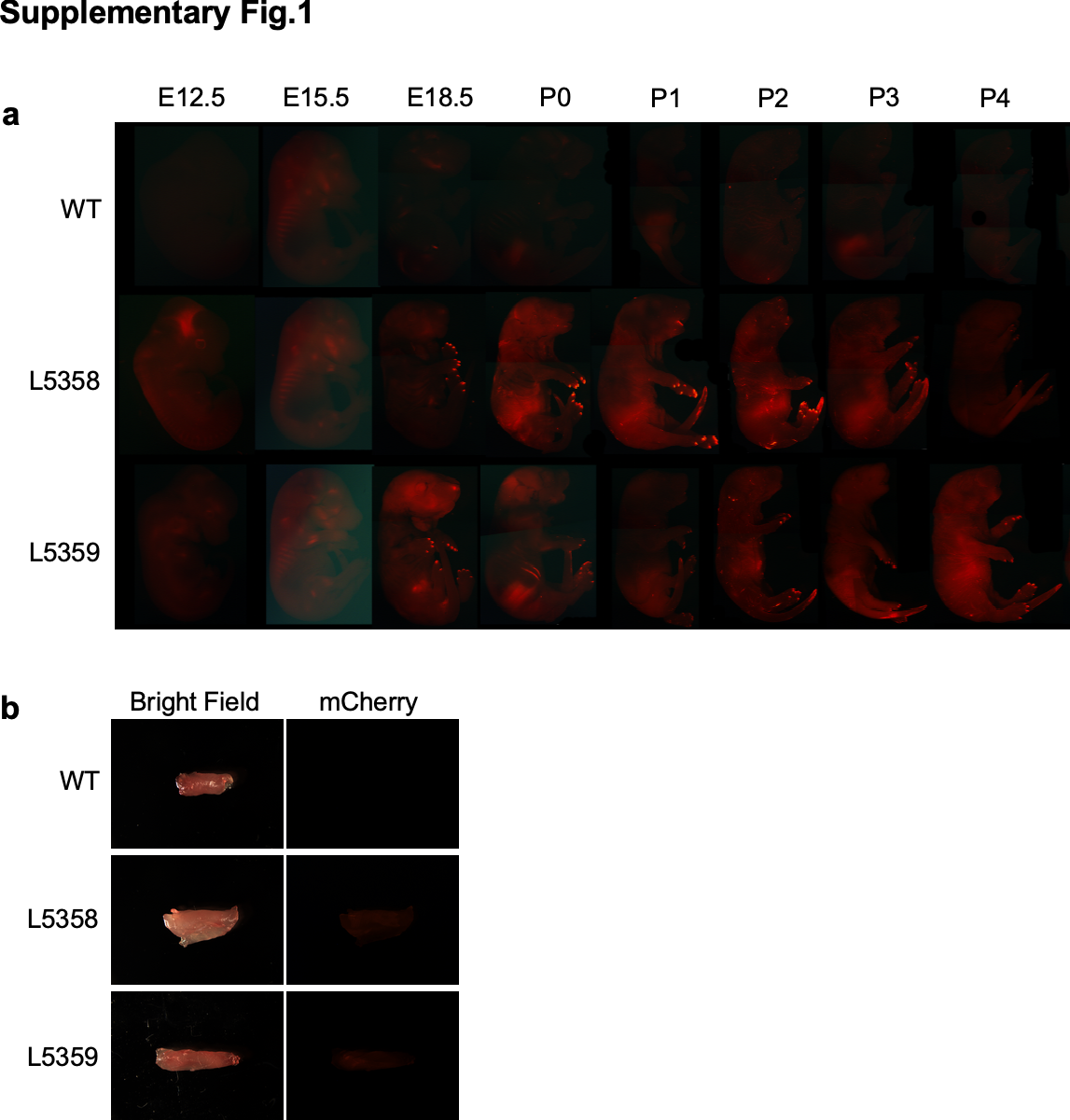

**Supplementary Fig. 1 | PEC7 enhancer transgenic assay. a,** mCherry expression in PEC7-HSP68-mCherry transgenic mice from E12.5 to P4 (wild type (WT), lines 5358 and 5339). **b,** mCherry expression in skeletal muscle from 10-week-old mice (WT, line 5358 and 5339).

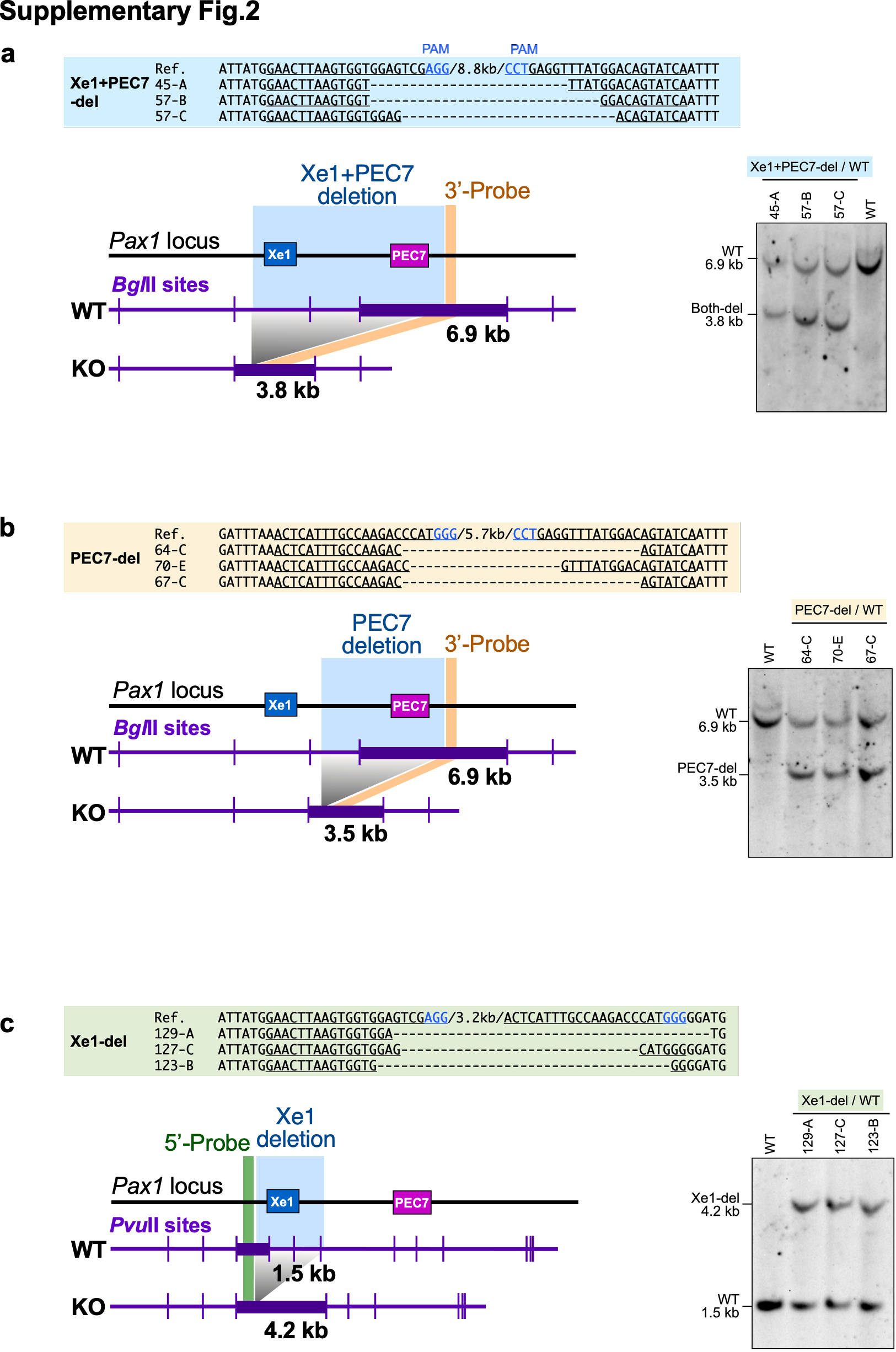

**Supplementary Fig. 2 | Sanger sequencing and Southern blot analyses of enhancer deleted alleles. a-c**, Sanger sequencing results for the deleted region. The PAM motif is depicted in blue font and gRNA sequences are underlined. Locations of the restriction enzyme sites (purple lines) in wild type (WT) and knockout (KO) loci. The deletion and probe locations are shown as orange and green lines respectively. Expected band sizes in WT and KO are indicated. Southern blot results of WT and KO mice with the estimated band sizes are shown.

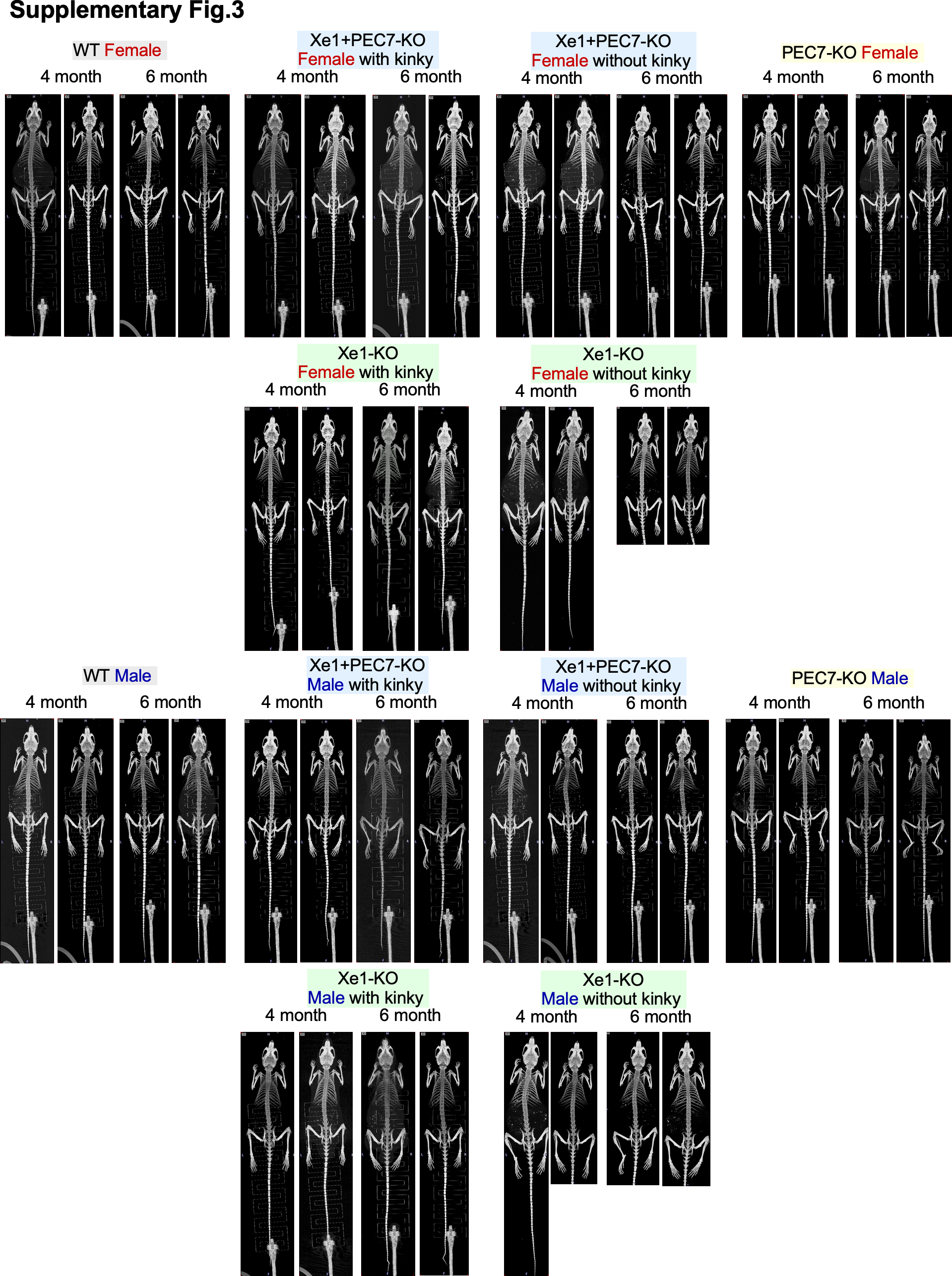

**Supplementary Fig. 3 | Whole-body skeletal structure analyzed by micro-CT.** Two mice in each genotype were analyzed at four months and six months of age. For Xe1 and Xe1+PEC7 mice with or without kinky tails were analyzed.

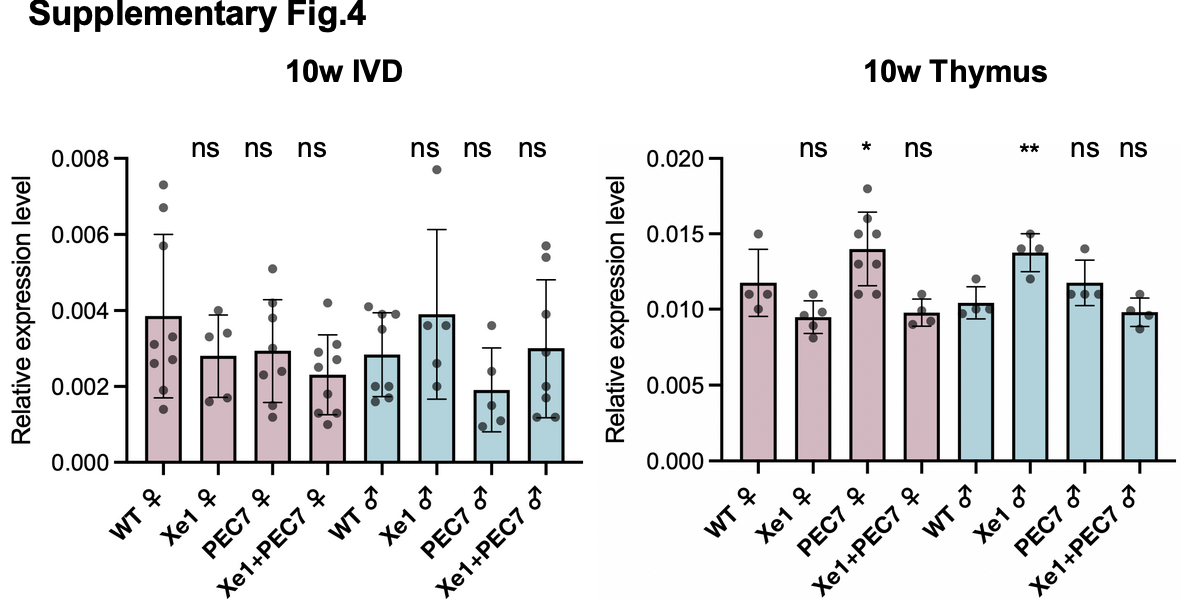

**Supplementary Fig. 4| Gene expression profiling for ten-week-old mouse tissues by qPCR.** Gene expression levels were dissected from ten-week-old mice as determined by qRT-PCR. Each value represents the ratio of gene expression to that of *β-Actin*, and values are mean ± standard deviation. Each dot represents one embryo. Statistical differences were determined using unpaired t test (**<0.01, *<0.05, ns, not signiﬁcant).

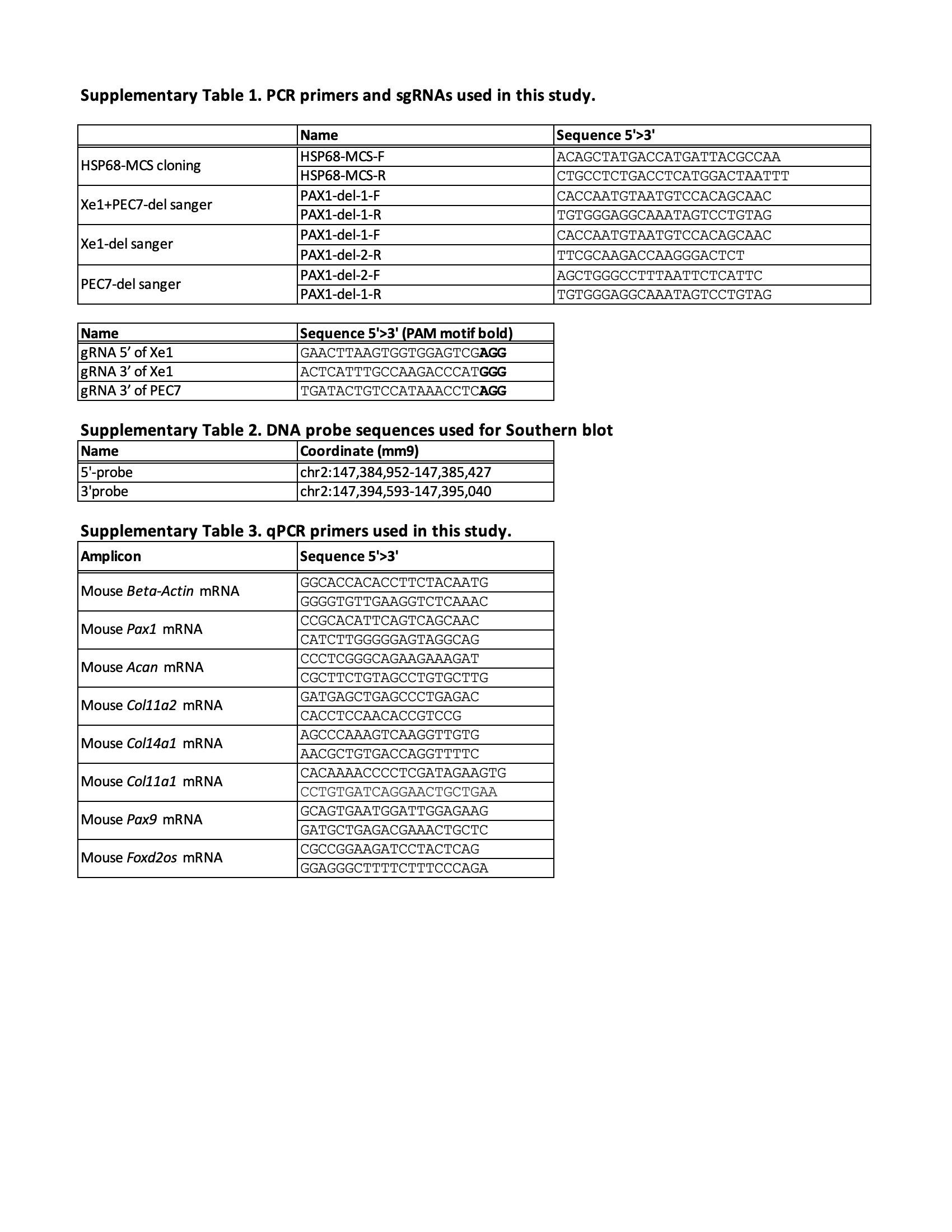
